# Trends in community- and nosocomial-acquired infections of carbapenem resistant Enterobacteriaceae (CRE), carbapenemase producing Enterobacteriaceae (CPE) and vancomycin resistant Enterococcus (VRE): a 10-year prospective observational study

**DOI:** 10.1101/2021.02.22.432249

**Authors:** Gloria Maritza Ubillus Arriola, William Araujo Banchon, Lilian Patiño Gabriel, Lenka Kolevic, María del Carmen Quispe Manco, José María Olivo Lopez, Armando Barrientos Achata, Maria Elena Revilla Velasquez, Donia Bouzid, Enrique Casalino

## Abstract

**Introduction:** CRE, CPE, and VRE are considered significant threats to public health.

**Aim:** To determine trends of nosocomial- and community-acquired infections.

**Methods:** A 10-year prospective observational non-interventional study was conducted. We used time-series analysis to evaluate trends in infections number.

**Findings:** Infection rate (%) were: CRE 2.48 (261/10,533), CPE 1.66 (175/10,533) and VRE 15.9 (121/761). We found diminishing trends for CRE (−19% [−31;−5], P=.03) and CPE (−22% [−30;−8], P=.04) but increasing trend for VRE (+48; [CI95% 34;75], P=.001). While we found decreasing trends for CRE and CPE in emergency (−71 [−122;−25], P=.001; −45 [−92;−27], P=.001) and hospitalization (−127 [−159; −85], P=.001; −56 [−98;−216], P=.01), we found increasing trends for VRE (+148 [113;192], P=.00001; +108[65;152], P=.003). Nosocomial-infections fell in CRE (−238 [−183;−316], P=.0001) and CPE (−163 [−96; −208], P=.001), but rose in VRE (+196 [151;242], P=.0001). We showed increasing trends in ambulatory and community-acquired infections in CRE (+134% [96;189]; P=.001; +77% [52;89]; P= .002), CPE (+288 [226;343]; P=.0001; +21% [−12;46]; P=.0.08) and VRE (+348 [295;458]; P=.0001; +66% [41;83]; P=.003). Direct admitted trends rose in all groups (CRE 16% [−8; 42]; P=.05), CPE 23% [−6; 48] (P=.05) and VRE (+241 [188; 301]; P=.0001).

**Conclusions:** We found a changing infection pattern with decreasing trends in in-hospital settings and nosocomial-acquired infections but increasing ambulatory and community-acquired infections. The observed increasing-trends in direct-admitted could be explained by community-onset infections diagnosed in the hospital. Our findings highlight the need to identify CRE/CPE/VRE community-acquired infections in ambulatory and in-hospital settings.

## Introduction

Carbapenem-resistant Enterobacteriaceae (CRE) are common pathogens causing severe infections and are responsible for increased mortality [1,2,3]. They also have significant clinical and socioeconomic consequences and are considered to pose a significant and severe emerging threat to public health [1–4]. Carbapenemase-producing Enterobacteriaceae (CPE) explain more of the observed rapid increase in CRE [5].

Enterococcus spp are inhabitants of the environment and normal intestinal microbiota. They may be pathogen opportunists, and they have been associated with urinary tract infections and sepsis, including bloodstream infections, abdominal infections, and endocarditis [6]. Vancomycin-resistant Enterococcus (VRE) has been reported since 1980. The mechanism is a result of changes in the formation of peptidoglycan that is coded by genotypes Van A to Van G [7]. Enterococcus spp is now recognized as a significant threat because of its increasingly frequent isolation in severe infections, its resistance to glycopeptides, and its multidrug resistance to last-resort antibiotics [6,7].

CRE, CPE, and VRE are considered emerging-highly resistant bacteria (eHRB) [8]. eHRB have been described mainly in hospital settings worldwide, increasing frequency among adults and children [1–7,9–11]. It has been recently reported that community-associated CRE/CPE rates range from 0% to 29.5% [11,12]. However, it is impossible to guarantee that they have not been acquired from previous healthcare exposure [12]. Community-acquired infections with VRE, however, have rarely been described outside the hospital setting [13].

Our goal is to determine trends in the number of CRE, CPE, and VRE in a pediatric population and determine their trends as a function of the hospital setting, type of infection acquisition, and hospital admission type.

## Materials and methods

### Study design

We conducted a 10-year prospective observational non-interventional study.

Study population, study period, and setting

All patients <15 years who attended the Instituto Nacional de Salud del Niño (INSN) with a clinically significant bacteriological sample between January 1, 2009, and December 31, 2018 (10 years) were included. The INSN is the national reference center for pediatrics in Peru, is in Lima-Peru.

### Definitions

All microbiological samples were prospectively recorded in a database. Patients’ clinical manifestations guide all samples. There is no screening for resistant bacteria carriage. Clinically significant samples were recorded prospectively by microbiologists and clinical physicians. All patients have a diagnosis code after each occurrence in all INSN settings. Based on the ICD-10 diagnoses, positive microbiological samples were associated with diagnostics. Variables included the type of hospital stay (ambulatory care, emergency, hospitalization); the type of infection (community-acquired or community-onset, hospital-acquired or nosocomial); and the type of admission (direct admission including through emergency room admission, transfer from another health care facility). The clinical files and previous visits at the INSN or other health structures were verified to define hospital-acquired and community-acquired origin by a group of experts (at least two authors, in case of disagreement, a third).

Primary endpoints were the number and rate per 100 diagnosed infections of CRE, CPE, and VRE.

### Characterization of resistance

eHRB included: i. Carbapenem-resistant Enterobacteriaceae (Escherichia coli, Klebsiella spp, Serratia marcesens, Enterobacter spp, and Proteus spp.); and ii. VRE (E. faecium and E. faecalis) [8].

We defined: i. Carbapenem-resistant Enterobacteriaceae (CRE) as non-sensitive to Carbapenem Enterobacteriaceae [1,2]; ii. Carbapenemase-producing Enterobacteriaceae (CPE): among CRE according to disc diameter interpretation algorithms and MIC [14,15]. Briefly, Enterobacteriaceae strains with reduced susceptibility to carbapenems (MIC ≤ 2/>8 mg/l for imipenem and meropenem or diameter zone for ertapenem <28 mm or for imipenem <24 mm, were tested on MH agar containing 250 mg/L cloxacillin, a diameter zone <15 mm for ticarcillin-clavulanate were considered CRE and then an in-house MALDI-ToF hydrolysis test was performed to detect the disappearance of the native carbapenem and/or the production of the carbapenem hydrolysis product after incubation of the tested strain with a carbapenem molecule [16,17]. These tests have been reported cost-effective with a negative predictive value of 100% and a sensitivity of 100% [14]. iii. VRE, according to standard interpreting antibiograms [18].

### Ethics

This study was approved by the INSN ethics committee on research. Ethics and clinical trial were registered in the Consejo nacional de ciencia, tecnología e innovación tecnológica (https://portal.concytec.gob.pe/).

### Data analysis

Data were aggregated by semester for rates evaluation or trimester for assessment of trends and ARIMA study. The numbers and percentages were first evaluated by Chi 2 or Fisher test and Student or Wilcoxon tests to compare qualitative and quantitative variables. Then, we used time series analysis (autoregressive integrated moving average [ARIMA] models) using the Box–Jenkins methodology. The method has been recently described in detail [19,20]. Moving averages provide a useful means of presenting time series data, highlighting any long-term trends while smoothing out any short-term fluctuations.

We evaluate intra- and inter-observer reliability for infection origin: nosocomial or community-acquired infections. All samples and files were reviewed by a pair of surveyors and classified blind (community-acquired or community-onset, nosocomial acquired). In the event of a discrepancy, a third investigator reviewed the file. The kappa (κ) statistic was used to denote agreement [21]. The significance threshold was 0.05. Missing data were not discarded. Statistical analyses were performed using Statistica software, version 12.5 (Statsoft/Tibco).

## Results

Fig 1 displays the study flowchart. Among 51,028 isolates, there were 17,625 clinically significant isolates of Enterobacteriaceae (34.5%), that include: K. pneumoniae (2,633 (14.9%)), K. oxytoca (552 (3.1%)), E. coli (13,527 (76.7%)), E. cloacae (329 (1.9%)), E. aerogenes (409 (2.3%)), S. marcescens (164 (0.9%)), and Citrobacter spp. (11 (0.06%)), P. mirabilis and P. Vulgaris (none). There were 911 isolates of Enterococcus spp. (1.8%), 687/911 (75.4%) were E. faecalis, and 224/911 E. faecium (24.6%). The main characteristics of the study population groups are presented in Table 1.

### Carbapenem resistance among selected Enterobacteriaceae with complete microbial evaluation

We found the following carbapenem resistance rates: K. pneumoniae 86/1831 (4.7%), K. oxytoca 19/130 (14.6%), E. coli 113/5797 (1.95%), E. cloacae 15/250 (6%), E. aerogenes 19/221 (8.52%), S. marcescens 9/126 (7.14%), and Citrobacter spp. 1/9 (11.1%).

### Vancomycin resistance among Enterococcus spp

We found the following vancomycin resistance rates: E. faecalis 26/566 (4.39%) and E. faecium 95/169 (56.2%)

### Overall infection rate

Infection rate (%) were for CRE 2.48 (261/10,533), CPE 1.66 (175/10,533) and VRE 15.9 (121/761).

### Validation of infection origin

After data and clinical evaluation, intra- and inter-observer’s kappa values for type of infection origin were 0.73 (CI95% 0.68-0.77) and 0.69 (0.64-0.74), respectively.

### CRE, CPE, and VRE number trends

We found during the study period significant increasing trends in the overall number of selected Enterobacteriaceae (+223% [CI95% 155; 288]; P< .000001) and Enterococcus spp. (+188% [CI95% 121; 249]; P< .000001).

As presented in Fig 2, we showed diminishing overall trends for CRE (−19% [CI95% −31; −5]; P= .03) and CPE (−22% [CI95% −30; −8]; P= .04), but an increasing trend for VRE (+48; [CI95% 34; 75]; P= .001).

We found increasing trends in ambulatory care for CRE (+134 [96; 189]; P=.0001) and CPE (+288 [226; 343]; P=.0001)), while we found decreasing trends for CRE and CPE in emergency ((−71 [−122; −25]; P=.001); (−45 [−92; −27]; P=.001)) and hospitalization ((−127 [−85; −159]; P=.001); (−56 [−21; 98]; P=.01)), respectively. VRE trends rose in ambulatory care (+272 [225; 358]; P=.0001), emergency ((+148 [113; 192] (P=.00001)) and hospitalization (108 [65; 152] (P=.003)). Whereas nosocomial infections trends fell for CRE (−238 [−183; −316]; P=.0001) and CPE (−163 [−96; −208]; P=.001), we found increasing trends in community-acquired infections for CRE (+77% [52; 89]; P= .002), CPE (+16% [−19; 46]; P=.0.05) and VRE (+114 [82; 139]; P =.001). Otherwise, we observed diminishing trends among transferred patients for CRE (−279% [−358; −192; P =.0001) and CPE (−152 [−216; −105]; P =.0001)) but increasing trends in direct admissions (CRE 16% [−8; 42]; P =.05), CPE 23% [−6; 48] (P =.05). For VRE, we found increasing trends in direct admission (+241 [188 to 301]; P =.0001) and transfer (+66% [41 to 88]; P=.005) (Fig 3).

## Discussion

Our study indicates a high ratio for CRE (2.48%), CPE (1.66), and VRE (15.9%) among clinically significant infections and for the whole eHRB (2.62%). During the 10-year study period, we found increasing trends for Enterobacteriaceae and Enterococcus spp. While the number of infections decreased for CRE and CPE in nosocomial, emergency, and hospitalization, their number of community-acquired and ambulatory settings increased significantly. VRE overall number, nosocomial and community-acquired, ambulatory and in-hospital settings, increased during the study period.

Previously reported CRE and VRE rate in healthcare settings are comprised between 0.25% and 9%, and 2% and 20%, respectively [1–7,9–11,22–25]. The CRE incidence rate in Europe and the USA is lower than 1% [26,27]. We found a CRE and VRE rates of 1.66% and 15.9% during the study period. Otherwise, we identified among CRE, 67% of carbapenemase-producing strains. Genes of carbapenem resistance have been reported in Latin America and Peru, mainly blaKPC and blaNDM [28,29]. Our study confirms that, as previously reported [30], the primary mechanism of Enterobacteriaceae resistance to carbapenems in Peru is carbapenemases production.

Community-acquired CRE, CPE, and VRE infections and carriage in children and adults have been previously reported [11–13,31,32]. A recent article indicates that although 33.3% of reviewed articles found no community-associated CRE, the remaining studies identified percentages ranging from 0.04% to 29.5% [12]. However, there have not differentiated community-associated and community-onset infections [12]. In the present study, we reviewed the patient’s files to define the infection origin, community- or nosocomial-acquired, with good results (kappa values for intra- and inter-observers were 0.73 (CI95% 0.68-0.77) and 0.69 (0.64-0.74)). We did not perform a systematic screening for eHRBs, but since all the patients included in the study have clinically significant samples, we have defined not the origin of carriage but the origin of documented infections.

In the present study, except for VRE, we found decreasing trends for CRE and CPE infection numbers in nosocomial, emergency and hospitalization settings. On the other hand, we found increasing trends in ambulatory care for CRE (+134 [96; 189]; P=.0001), CPE (+288 [226; 343]; P=.0001) and VRE (+272 [225; 358]; P=.0001). Evermore, we found increasing trends in community-acquired CRE (+77% [52; 89]; P= .002), CPE (+21% [−12; 46]; P=.0.08) and VRE (+114 [82; 139]; P=.001). Our results indicate a changing pattern in eHRB origin, with less frequent nosocomial infections but an increasing number of community-acquired infections and ambulatory care.

Interestingly, we observed an epidemic episode of CRE, CPE, and VRE during the last semesters of the study period, mainly among directly admitted patients (Fig 2). Some isolated strains were identified in the first bacteriological sample set and were responsible for the admission diagnosis. We believe that the increase in the number of infections in direct-admitted patients resulted from the increased number of eHRB in the community in patients without previous contact with the health care system. Even if carbapenemase-producing Enterobacteriaceae were usually associated with hospital dissemination [33], it has been recently reported that strains-producing carbapenemases dissemination occurs outside the hospital [34], and that CRE/CPE and VRE were frequently identified in hospital and community wastewater [35], which may reflect the dissemination of these strains, in particular VREs, in the community. To our knowledge, our study is the first one to evaluate the community-onset of clinically significant infections in different care settings and their trends over an extended observation period, which allows us to demonstrate an increased frequency of community-onset infections, some of them diagnosed during a hospital stay.

Our study has some limitations and strengths: first, the participants were enrolled from one hospital, but INSN is a national pediatric reference center and receives patients from all Lima agglomeration and Peru; second, patients have not been systematically screened for eHRB ad admission. However, al eHRB infection episodes were investigated with a rigorous and validated method to define infection origin.

## Conclusions

In conclusion, our study confirms the high frequency of eHRB in the pediatric population and a changing pattern in eHRB infections with increasing trends in the ambulatory setting and for community-acquired infections. Our study demonstrates that eHRB, regardless of whether it remains a nosocomial and care-related problem, should also be considered as a problem in ambulatory care and in-hospital settings, even in patients without any documented previous contact with the healthcare environment. Finally, our findings indicate and highlight the need for new research lines to determine their frequency and risk factors and define the relevance of new screening and isolation strategies in ambulatory and hospital settings. As recently suggested, efforts should be placed on controlling the spread of eHRB in the community [36].

## Acknowledgments

We thank all the INSN healthcare workers who participated in this study.

## Supporting information

**S1 Fig.**
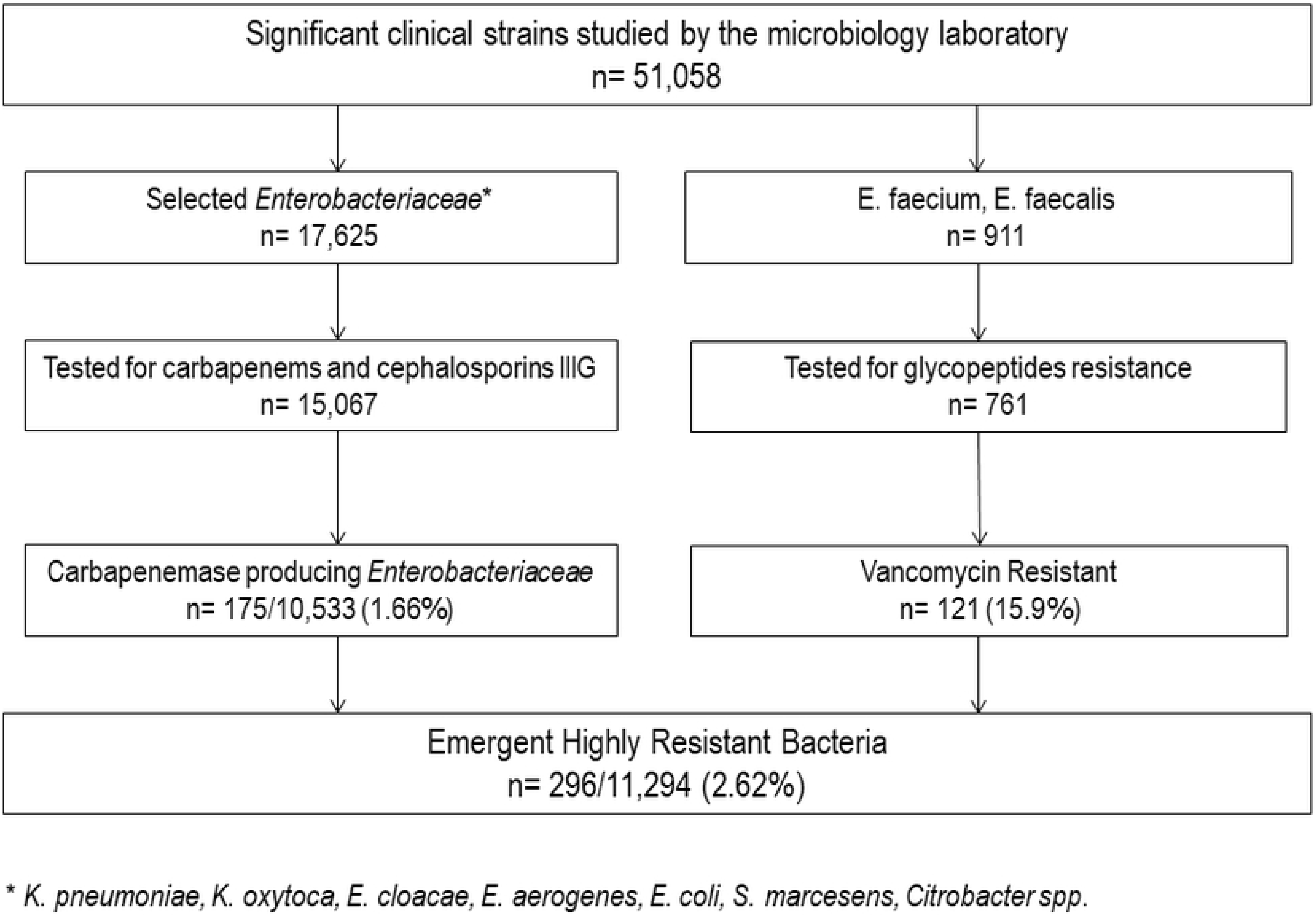
Study flowchart.

**S2 Fig.**
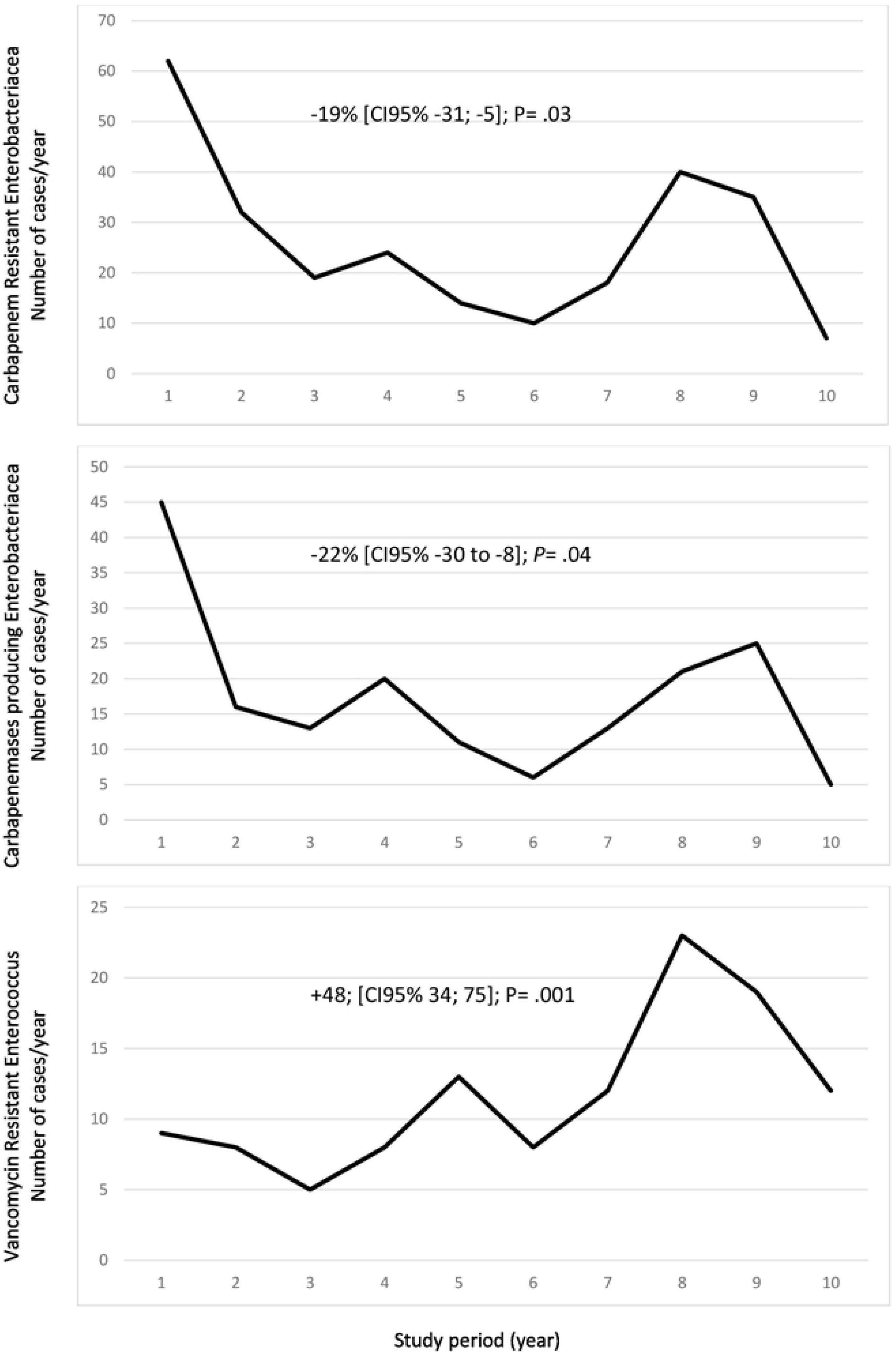
Overall trends of Carbapenem resistant Enterobacteriaceae, Carbapenemase producing Enterobacteriaceae and Vancomycin resistant Enterococcus during the 10 years study period.

**S3 Fig.**
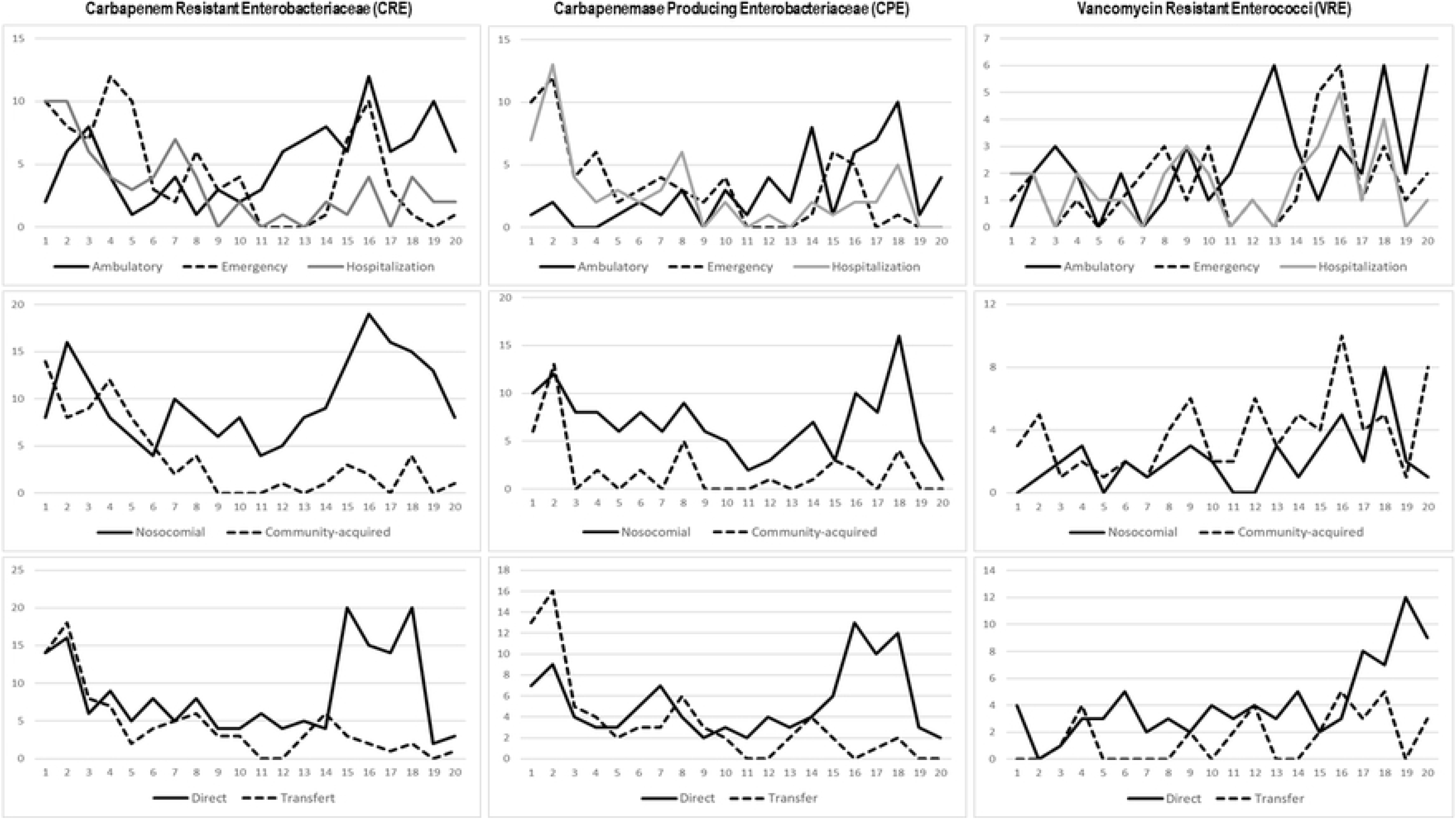
Trends of Carbapenem resistant Enterobacteriaceae, Carbapenemase producing Enterobacteriaceae and Vancomycin resistant Enterococcus as a function of infection characteristics.

S1 Table. Main characteristics of selected Enterobacteriacea and Enterococcus strains.

